# Compressed sensing expands the multiplexity of imaging mass cytometry

**DOI:** 10.1101/2023.11.06.565119

**Authors:** Tsuyoshi Hosogane, Leonor Schubert Santana, Nils Eling, Holger Moch, Bernd Bodenmiller

## Abstract

The multiplexity of current antibody-based imaging is limited by the number of reporters that can be detected simultaneously. Compressed sensing can be used to recover high-dimensional information from low-dimensional measurements when the data has a structure that allows sparse representation. Previously, in composite in situ imaging (CISI) of transcriptomic data, compressed sensing leveraged the gene co-regulation structure that allows sparse representation and recovered spatial expression of 37 RNA species with the measurement of 11 fluorescent channels. Here, we extended the compressed sensing framework to protein expression data measured by imaging mass cytometry (IMC). CISI-IMC accurately recovered spatial expression of 16 proteins from the images of 8 composite channels, which in effect expanded the current multiplexity limit of IMC by 8 channels. With this ratio, up to 80 protein markers could be compressed into currently available 40 isotope channels. Training the CISI-IMC framework using data collected on tissues from various locations in the human body enabled the decompression of composite data from a wide range of tissue types. Our work laid the foundation for much higher plex protein imaging by using CISI.

## Introduction

Antibody-based protein imaging allows profiling the spatial distribution of cell phenotypes across tissues, therefore forming a crucial basis to understand the pathology of diseases. However, detecting biomolecules in standard immunofluorescence imaging is limited by the number of available fluorescent channels, typically five, due to spectral overlap.

This multiplexity limitation has been overcome by performing iterative staining and de-staining cycles using few fluorescently-labeled antibodies in each cycle. In most protein imaging methods, each channel in each round of staining corresponds to a specific marker^1–6^. The iterative methods such as CODEX, 4i, and CyCIF can achieve up to 60-plex imaging^2,3,7^. Higher multiplexity has been achieved using combinatorial barcoding, where the sequence of channels over the rounds for each target molecule creates a unique barcode. For example, CosMx, which employs combinatorial barcoding, has been used to image 108 proteins in a tissue sample^8^. Limitations of these methods are autofluorescence and tissue damage caused by repeated staining cycles. In addition, the combinatorial barcoding approach requires the single molecule detection of each target molecule for accurate decoding and thus is confounded by molecular crowding. Even higher multiplexity can be achieved by sequencing-based approaches, in which antibodies are labeled with a DNA tag that carries a barcode that can be sequenced to identify the location on the slide^9^. For example, 300-plex imaging was demonstrated with spatial CITE-seq^10^. Sequencing-based methods do not suffer from autofluorescence and do not require repeated staining cycles, but currently the resolution is too low to allow single-cell analyses (25 μm for spatial CITE-seq).

In contrast to the fluorescent-based and sequencing-based methods, mass spectrometry-based imaging methods such as imaging mass cytometry (IMC) and multiplex ion beam imaging (MIBI) enables up to 40-plex imaging without sequential staining cycles at a spatial resolution of 1 μm or higher^11,12^. In IMC, antibodies are tagged with metal isotopes that are simultaneously detected by laser ablation and subsequent mass spectrometry. The multiplexity of IMC is limited by the number of available channels that are used to detect individual metal isotopes. Since IMC is a destructive imaging method, iterative staining cannot be used to expand the multiplexity. Combinatorial barcoding could be implemented by labeling each antibody with a unique combination of metal isotopes, but the current IMC resolution of 1 μm is too low to achieve single-molecule detection that is required for accurate decoding. Expanding IMC beyond 40-plex would be possible with a method that achieves the decoding of combinatorial barcoding without the need for single-molecule detection.

In the RNA imaging field, composite in situ imaging (CISI) achieves such a decoding of combinatorial barcodes without single-molecule detection by using prior knowledge of transcriptome profiles^13,14^. Leveraging the biological principle of gene co-regulation, CISI constructs a dictionary of gene expression modules from training data, which enables the decoding of the combinatorial barcode imaging data into the spatial expression data of individual genes. By using a dictionary of gene expression modules obtained by training on single-cell RNA sequencing (scRNA-seq) data, CISI accurately recovered the spatial abundance of 37 RNA species from the combinatorial barcode imaging data of only 11 channels (3 and 2/3 rounds of iterative staining, 3 colors each round)^14^. Importantly, CISI decoding can be performed on the single-cell level (or pixel-level when using an autoencoder), and therefore only requires resolution high enough for single cell analysis.

Here, we implemented CISI for IMC to expand its multiplexity over the number of available isotope channels. We reasoned that a protein expression dictionary for IMC data could be constructed since proteins are also co-regulated. Further, the current IMC resolution of 1 μm is sufficient to perform CISI on the single-cell level. We performed 16-plex standard IMC imaging of multiple tissues using antibodies that mainly identify immune and stromal cells for training data and constructed a protein expression dictionary. We then implemented combinatorial barcoding by mixing 16 antibodies each labeled with unique combination of 8 metal isotopes. We demonstrated that the obtained 8-channel CISI-IMC images for five different tumor tissues were accurately decoded back to 16-plex spatial single-cell protein expression data using our optimized CISI algorithm.

## Results

### CISI-IMC principles

Using CISI applied in IMC, we recover high-dimensional single-cell protein expression data from low-dimensional single-cell composite data obtained by combinatorial barcoding of antibodies. To obtain low-dimensional single-cell composite data in IMC, antibodies targeting *p* proteins are labeled with unique combinations of *m* isotopes, where *m* < *p* (Fig. 1a). The barcoding matrix *A* describes the combination of isotopes labeled for each antibody. By measuring a tissue stained with the barcoded antibodies by IMC, single-cell protein expression of *p* proteins is linearly combined and compressed into single-cell composite data with *m* channels (Fig. 1a). The matrix *X* contains single-cell protein expression profiles and the matrix *Y* describes the single-cell intensity profile of composite isotope channels.

**Fig. 1.**
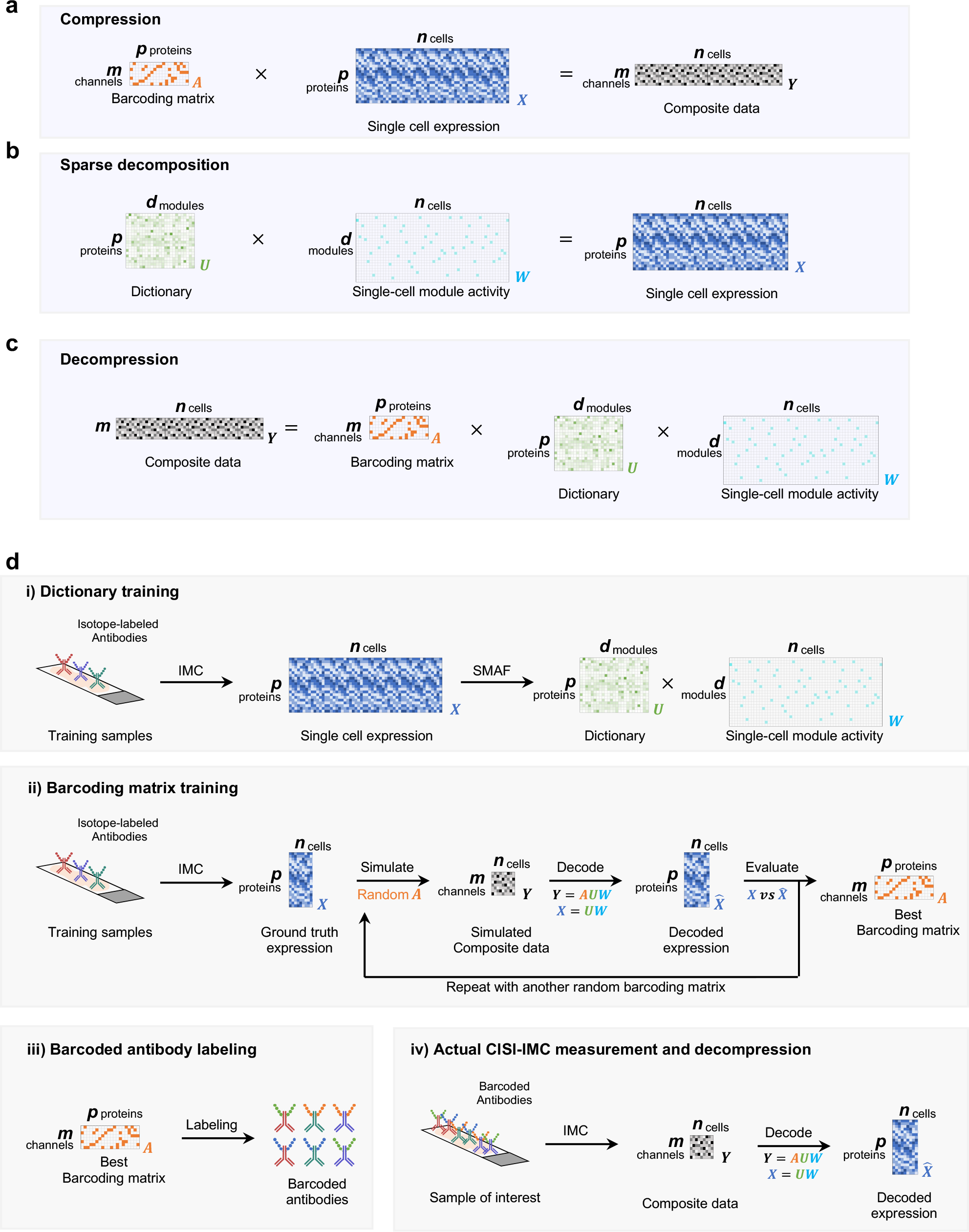
CISI-IMC workflow. **a**, Schematic of the compression process using CISI-IMC. The barcoding matrix encodes the scheme for barcoding of *p* antibodies with unique combinations of *m* composite channels. By selecting an *m* smaller than *p*, single-cell protein expression data of *p* proteins can be compressed into single-cell composite data of *m* composite channels. **b**, Schematic of the sparse decomposition using CISI-IMC. To enable the decompression, single-cell protein expression data of *p* proteins is decomposed into a dictionary of *d* protein-expression modules and single-cell module activity with sparse activation of the modules for each cell. **c**, Schematic of decompression using CISI-IMC. By combining the equations for compression and sparse decomposition, composite data can be decomposed into a barcoding matrix, a dictionary, and a single-cell module activity matrix. Decompression is performed by estimating the sparse single-cell module activity using the known barcoding matrix and the dictionary. **d**, CISI-IMC experimental workflow. (i) The dictionary is trained using single-cell IMC data from a training dataset and the SMAF algorithm. (ii) The barcoding matrix is trained on a portion of the training dataset by simulating the single-cell composite data with randomly generated barcoding matrices. Simulated single-cell composite data are then decompressed back to single-cell protein expression data, which are compared to the original single-cell protein expression data, and the barcoding matrix with the best decompressing performance is selected for the next steps. (iii) Antibodies are labeled with the specific combinations of metal isotopes according to the selected barcoding matrix. (iv) Tissues of interest are stained with the barcoded antibodies and imaged with IMC. Obtained composite images are segmented into single-cell composite data that is decompressed back to single-cell protein expression data using the pre-trained dictionary.

To aid the decompression, single-cell protein expression *X* is decomposed into two matrices; one is a dictionary of protein-expression modules and the other is single-cell module activity (Fig. 1b). The protein-expression module describes a pattern of protein expression, and the dictionary stores different protein-expression modules. For each cell, a few modules are selected from the dictionary and linearly combined to approximate the single-cell protein expression. The dictionary *U* describes protein expression patterns for each module, and the single-cell module activity *W* describes which module is active for each cell (Fig. 1b). In CISI, the dictionary *U* is pre-defined using a training dataset. The decompression task is facilitated by the sparsity constraint of *W* which defines that only a few modules can be active for each cell.

Obtained single-cell composite data *Y* is decompressed back to individual protein expression data by decomposing *Y* into the barcoding matrix *A*, the dictionary *U*, and single-cell module activity *W*, using the previous two equations from compression and decomposition (Fig. 1c). Since the barcoding matrix *A* and the dictionary *U* are known and the single-cell module activity *W* has to be sparse, it becomes mathematically possible to estimate *W*, and to reconstruct single-cell protein expression *X* by *X* = *UW*.

### CISI-IMC workflow

The CISI-IMC workflow consists of four major steps: dictionary training, barcoding matrix training, barcoded antibody labeling, and CISI-IMC measurement and decompression into individual protein expression (Fig. 1d). In the dictionary training step, single-cell protein expression data obtained from IMC measurements of various tissue samples were used as a training dataset. The sparse module activity factorization (SMAF) algorithm was used to construct a dictionary that well-approximates single-cell protein expression with sparse single-cell module activity. In the following barcoding matrix training step, another portion of the training dataset was used to simulate the single-cell composite data using randomly generated barcoding matrices. Simulated single-cell composite data were then decompressed into single-cell protein expression data that was compared against the original single-cell protein expression data. The barcoding matrix with the best decompressing performance was selected for the next steps. In the barcoded antibody synthesis step, antibodies were labeled with metal isotopes according to the selected barcoding matrix. Finally, CISI-IMC measurements were carried out: Tissues of interest were stained with the barcoded antibody mix and imaged with IMC, and obtained composite images were segmented into single-cell composite data, which were decompressed into single-cell protein expression data using the pre-trained dictionary.

### SMAF optimization for CISI-IMC

We first examined whether the SMAF algorithm from the CISI framework^14^ can be applied to protein expression data as generated by IMC. Here we used protein expression data across six different tumors and three different healthy tissues from various organs. We developed a panel of 16 protein markers to detect various cell types that could be found across tissues (Supplementary Table 1). SMAF decomposes single-cell protein expression data *X* into a dictionary of protein expression modules *U* and a single-cell module activity matrix *W*, while enforcing sparsity for both *U* and *W* within the provided error tolerance (that is, the distance between *X* and *UW*). *U* and *W* were calculated by iteratively finding the sparse solution for one while fixing the other (Extended Data Fig. 1a). The Lasso algorithm was used to calculate *U* from a fixed *W*. Lasso and orthogonal matching pursuit (OMP) were tested for calculating *W* from a fixed *U*. Lasso calculates the sparsest solution within the provided error tolerance coefficient *lda*. OMP finds the best fit solution within the provided sparsity *k* (that is, up to *k* modules per cell are non-zero). We found that 100 iterations were sufficient to obtain stable solutions for most of the conditions tested (Extended Data Fig. 1b, c). After 100 iterations, SMAF was able to find sparse *U* and *W* within the error tolerance (Extended Data Fig. 1d-k). As expected, a looser error tolerance coefficient for calculating *U* (larger *ldaU*) resulted in sparser *U* and reduced the total number of modules in *U* (Extended Data Fig. 1d, e, h, i). In addition, reducing the error for calculating *W* (smaller *ldaW* or larger *k*) also resulted in sparser *U*, likely because denser *W* allows sparser *U* within the same error tolerance in total (Extended Data Fig. 1d, f, h, j). Separating cells in *X* into blocks based on the vector size when calculating *W* (specified by *Num*_*blocks*_*W*) slightly reduced the sparsity of *U* (Extended Data Fig. 1d, e). Although SMAF produces slightly different *U* values over experimental replicates, the difference was minimal when the Lasso error tolerance coefficients (that is, *ldaW* and *ldaU*) were not overly stringent (Extended Data Fig. 2).

### Simulating CISI-IMC for optimizing the training protocol

Next, we simulated the single-cell composite data *Y* using randomly generated barcoding matrices *A* on the single-cell training data *X*. Since each composite channel in *Y* is a linear sum of proteins defined by *A*, we simply multiplied *X* by *A* to obtain simulated *Y*. The obtained *Y*was decompressed back into *X* using Lasso on *Y* with *AU* (that is, *Y* = *Lasso*(*X, AU*)). Decompression performance was evaluated for each protein by calculating the Pearson’s correlation coefficient between the decompressed *X* and the original *X*. Each protein in a randomly generated *A* was restricted to have only one or two composite channel entries to minimize the complexity during the antibody labeling. Since the number of non-zero composite channels per protein in *A* is equivalent to how many different isotopes are used to label a certain antibody, the imposed restriction guarantees that each antibody was labeled with no more than two isotopes. We observed that denser *A* generally resulted in better decompression performance and that increasing the maximum composite channels per protein to three or four did not improve the performance for the tested compression rate of 16 proteins into 8 composite channels (Extended Data Fig. 3). Therefore, we used no more than two isotope labels per protein and the number of proteins with only one composite channel in randomly generated barcoding matrices was restricted to no more than four to enforce the denser *A*.

Normalizing proteins in *X*, that is, protein-wise scaling to enforce each protein data in *X* to have the same vector size, when simulating *Y* improved the decompression performance (Extended Data Fig. 4a, b). This was expected since *X* was also normalized for proteins when calculating *U* via SMAF, thus the values of proteins in each module, which would be linearly combined to approximate *X* for decompression, was based on the normalized intensity. However, normalizing proteins in *X* in actual CISI-IMC staining is not straightforward. Considering that there exist positive and negative cell populations for each protein marker, we need to account for two main factors that affect the norm of *X*: the signal intensity of positive cell population and the density of positive cell population. In other words, a protein expressed only on a rare cell population should have higher signal intensity than a protein expressed on a common cell type in order to accurately normalize proteins in *X*. Although the signal intensity for each marker can be adjusted by antibody titration, the abundance of a cell type is dependent on the tissue sampled. Adjusting signal intensity based on cell type abundance observed in a training dataset hinders the applicability of the method to various tissue types. Therefore, we aimed for uniform signal intensity for each marker-positive cell population, which should minimize the effect of varying cell type abundance. We prepared another training data *X* using titrated antibodies on the same tissues as used in the original training dataset. Simulating decompression using this training data indeed showed improved performance (Extended Data Fig. 4c, d).

### Training a barcoding matrix and a dictionary for CISI-IMC

To finalize the dictionary *U*, we calculated *U*s with 12 different SMAF parameter conditions on 75% of the training data using titrated antibodies and simulated *Y*. On the rest of the trainings data, we evaluated decompression performance using 200 randomly generated *A*s (Extended Data Fig. 5a). In addition, performance stability across tissues was assessed by evaluating the decompression performance for each tissue using a *U* calculated on the training data excluding the data from the tissue of interest (Extended Data Fig. 5b). The final SMAF parameters were selected based on this experiment and on the performance stability across different normalization weights for training data, which simulates the variability in antibody titration (Extended Data Fig. 5c). The final *U* was obtained using these finalized SMAF parameters (*algorithm*_*for*_*W*: *Lasso, ldaU*:0.02, *ldaW*:0.02, *Num*_*blocks*_*W*: 1) (Extended Data Fig. 5d).

To finalize the barcoding matrix *A*, we generated random *A*s barcoding 16 proteins in 8 composite channels, with each protein maximally using 2 channels. We simulated *Y* and observed decompression performance using 2,000 randomly generated *A*s on 25% of training data using *U* calculated with the finalized SMAF parameters on the rest of the training data. We performed the simulation four-fold and selected the top 50 *A*s from each based on the Pearson’s correlation for the worst performing protein, which resulted in 200 candidates from 8,000 *A*s tested. Finally, we performed another set of simulations fixing the 200 *A* candidates and selected the best *A* based on the highest minimum protein correlation averaged over the four-fold simulations. For the best *A*, the simulated average protein correlation coefficient was 0.924 and minimum protein correlation coefficient was 0.864. We also tested the

*A*s with 7 and 9 composite channels. As expected, increasing the number of composite channels improved the decoding results (Extended Data Fig. 6a). We selected 8 composite channels to balance performance and efficiency of the decompression (Extended Data Fig. 6b).

### Actual CISI-IMC experiment compressing 16 proteins into 8 channels

Barcoded antibodies compressing 16 proteins into 8 composite channels were labeled according to the finalized *A*. All unique antibody-isotope pairs were conjugated and titrated separately and pooled together as a barcoded antibody mix. To evaluate the decompression performance of CISI-IMC experimentally, we stained five tumor tissues with the barcoded antibody mix together with a ground-truth antibody mix, which were antibodies each conjugated to a single isotope. Hence, the obtained evaluation dataset of five tumor tissues contained single-cell composite data *Y* with matching ground-truth single-cell protein expression *X*. The ground-truth *X* was used to evaluate the accuracy of decompressed 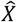 and to produce a simulated *Y* to assess the accuracy of the simulation. For the decompression of the evaluation dataset, we used the dictionary *U* calculated from the training dataset, because we hypothesized that *U* from a diverse training dataset would be sufficient for the decompression of different tissues provided that the expression patterns for the selected 16 cell type protein markers were stable across tissues.

The decompression algorithm from the previous publication^14^ was able to recover individual protein expression with the protein average Pearson’s correlation coefficient of 0.746 between decompressed 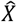 and ground truth *X* (Extended Data Fig. 7a, b). However, some proteins such as MPO were not accurately decompressed (Pearson correlation coefficient of 0.255). These correlation values were lower than the simulated values in the training dataset (the average protein correlation coefficient was 0.924 and the minimum protein correlation coefficient was 0.864). We hypothesized that the inaccurate decompression arises due to two reasons. First, the dictionary *U* calculated from the training dataset might not have been broadly applicable enough to decompress the evaluation dataset. Second, the actual *Y* obtained from barcoded antibodies might not have been accurately simulated during the training steps.

To test the first hypothesis, we prepared another dictionary *U* calculated on the evaluation dataset itself. Decompression performance of the evaluation dataset did not improve substantially when using the *U* from the evaluation dataset itself, suggesting that the weak performance for a few proteins was not caused by the used training dataset (Extended Data Fig. 7a, b).

To test the second hypothesis, we simulated *Y* using ground truth *X* from the evaluation dataset and compared it against the actual *Y*. We observed that the composite channel containing MPO had a lower correlation coefficient (0.646) between the actual *Y* and the simulated *Y* compared to the other composite channels (Extended Data Fig. 7c). In addition, proteins that poorly performed in the actual decompression, including MPO, were decompressed more accurately when using the simulated *Y*, suggesting that the reduced performance of a few proteins is at least partially due to the discrepancy between the actual *Y* and the simulated *Y* (Extended Data Fig. 7a, b). We further speculated that the actual *Y* was not accurately simulated because the weights in the barcoding matrix *A* were not reflecting the actual weight of signal intensity of each antibody-isotope conjugate. Since evaluating signal intensities of all antibody-isotope conjugates used in the barcoding matrix *A* for each tissue is not practical for highly multiplexed CISI-IMC, we developed an algorithm to reweight *A* post-hoc. By iteratively calculating the reweighted barcoding matrix *Â* and the module expression 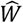 during the decompression until the decomposition error (that is, the distance between *Y* and 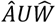) was minimized, we obtained a reweighted *Â* that can more accurately simulate the actual *Y* (Extended Data Fig. 7c). Using the reweighted Â, the decompression performance of actual CISI-IMC in the evaluation dataset improved to a protein average correlation coefficient of 0.800 and a minimum protein correlation coefficient of 0.507 (Fig. 2a, b). To evaluate if the decompressed single-cell expression data can be used for downstream analyses, we clustered cells using Phenograph on either ground truth *X* or decompressed 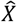. Each cluster was manually annotated into specific cell types based on known marker expression. All cell types assigned using ground truth *X* could also be assigned using decompressed 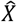 (Extended Data Fig. 8a, b) and the decompressed 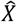 well-preserved the separation between cell types in UMAP embedding (Extended Data Fig. 8c, d). Finally, we assessed if the decompressed 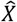 can accurately classify cell types by comparing the annotated cell types using decompressed 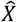 against those using ground truth *X* (Fig. 2c). All cell types were well recovered using decompressed 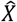 with average F1 score of 0.564 (Fig. 2d).

**Fig. 2.**
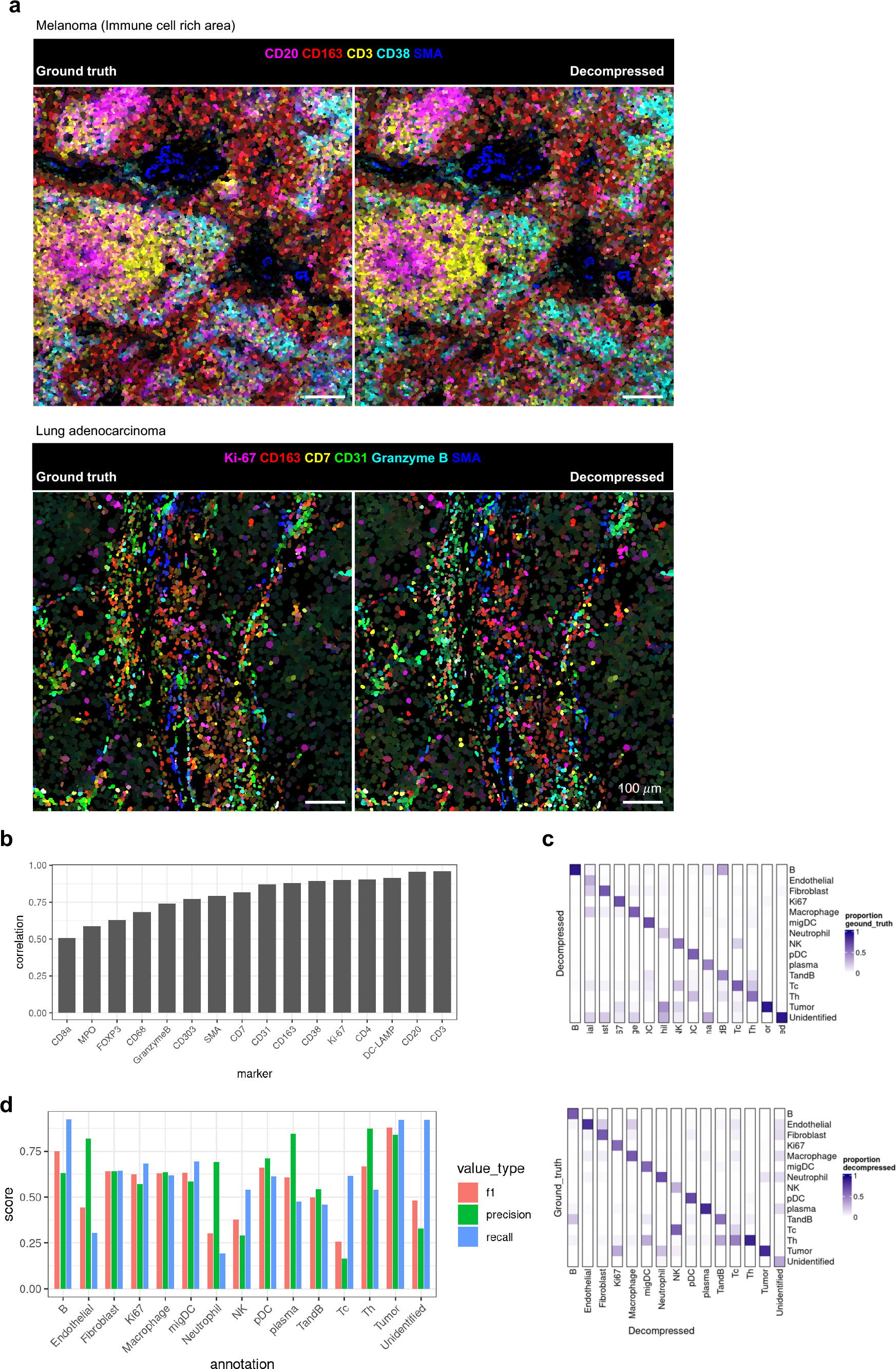
Performance of CISI-IMC on tumor tissues compressing 16-plex proteins into 8-plex composite channels. **a**, Example images of ground-truth cell image and decompressed cell image from immune-cell rich region of melanoma (top) and lung adenocarcinoma (bottom). Displayed protein markers are indicated on top of the images. Scale bars are 100 μm. **b**, Pearson correlations between decompressed and ground-truth single-cell expression data for each protein. **c, (top)** Proportions of annotated cell types based on decompressed data displayed for each annotated cell type based on ground truth data. **(bottom)** Proportions of annotated cell types based on ground truth data displayed for each annotated cell type based on decompressed data. **d**, F1-score, Precision, and Recall for each annotated cell type.

## Discussion

Here we demonstrated that CISI can extend the multiplexity of IMC beyond the number of available isotope channels. The SMAF algorithm was used to create a dictionary of protein expression modules based on a training dataset, and single-cell expression data for 16 proteins were accurately recovered from the measurement of 8 composite channels. In effect, this expands the multiplexity of IMC by 8 channels. We also demonstrated that use of a training dataset based on images of different tumor and healthy tissues resulted in a dictionary universal to different tissue types.

The reduced decompression performance for some markers was attributed to the discrepancy between the actual composite data and the simulated composite data rather than to the difference of protein expression pattern in the training dataset and the evaluation dataset. Previous application of CISI to RNA imaging has shown that transcripts that are rarely expressed and lowly abundant transcripts were less accurately recovered. In this work, we did not increase the intensity for rare and low-abundance proteins by increasing antibody concentration, since expression levels differ between tissues. However, we selected antibody concentrations that resulted in similar signal intensities for cells positive for each protein. In addition, reweighting the barcoding matrix during the decompression largely improved the performance for the poorly recovered proteins.

To implement CISI-IMC, standard IMC data (or other highly-multiplexed imaging data) can be used to simulate CISI and predict the decompression performance for the selected proteins. After obtaining the dictionary and the barcoding matrix from training on the standard IMC data, antibodies can be barcoded and the tissue of interest can be stained and imaged. Obtained composite data can be decompressed back to individual protein expression data using the pre-trained dictionary.

Increasing the number of composite channels resulted in improved decompression performance in our simulation. Therefore, depending on the accuracy of decompression required for a project, the compression ratio can be adjusted. In this manuscript, we used two-fold compression from 16 proteins into 8 composite channels. With this ratio, up to 80 protein markers can be compressed into currently available 40 isotope channels.

We reason that cell-level decompression will be sufficient for most multiplexed IMC protein analyses, since standard IMC data are almost always aggregated within each cell mask and spatial analyses are performed on the single-cell level. In previous work, pixel-level decompression using deep learning models has been demonstrated^14,15^, and this strategy could theoretically be implemented for IMC. In cases where subcellular localization of protein expression is of interest, pixel-level decompression would be beneficial.

In summary, CISI-IMC is a highly multiplexed imaging approach that allows simultaneous detection of 48 protein markers and potentially up to 80-plex or even higher number of proteins. Current multiplexed protein imaging techniques with single-cell resolution can achieve approximately 60-plex imaging, but all rely on iterative imaging cycles, which requires complex image processing steps. IMC and MIBI are the exceptions that do not require iterative imaging cycles, but their multiplexities are limited to about 40 plex due to the number of available isotope channels. CISI-IMC retains the non-cyclic nature of IMC but expands the multiplexity over the limited number of isotope channels.

## Methods

### FFPE sections of human tumor samples for training and evaluation datasets

The training dataset was derived from six human tumor tissues (lung adenocarcinoma, lung squamous cell carcinoma, colon adenocarcinoma, invasive lobular breast carcinoma, and two breast cancer) and three human healthy tissues (tonsil, lung, and appendix). The evaluation dataset was derived from five human tumor tissues (lung adenocarcinoma, lung squamous carcinoma, clear cell renal cell carcinoma, melanoma, and ovarian cancer). FFPE sections were prepared at University Hospital Zurich. All the FFPE sections were kept at room temperature for short-term storage or at -20 °C for long-term storage. Use of samples received from University Hospital Zurich was approved by the Ethikkommission Kanton Zürich (KEK-ZH-Nr. 2014-0604, 100TO).

### Antibody labelling with metal isotope

Isotope-labeled antibodies were used for IMC staining of training samples and evaluation samples. Antibodies were labeled with an isotope using the MaxPar X8 Antibody labelling kit (Fluidigm) according to the protocol supplied by the manufacturer. First, chelation was completed by incubating MaxPar X8 polymer, which carries terminal maleimide functionality and multiple chelators for lanthanide ions, in 2.5 mM lanthanide chloride solution (Fluidigm) at 37 °C for 30 min. The product was purified into C-buffer, provided with the MaxPar X8 Antibody labeling kit, using 0.5-ml, 3-kDa Amicon Ultra Filters (Millipore). In parallel, antibody was partially reduced in 0.8 mM TCEP at 37 °C for 30 min and was purified into C-buffer using 0.5-ml, 50-kDa Amicon Ultra Filters. Isotope-loaded polymer was added into partially reduced antibody and was incubated at 37 °C for 90 min. Conjugated product was purified over 0.5-ml, 50-kDa Amicon Ultra Filters. For CISI-IMC, each pair of isotope and antibody was separately conjugated and purified. Antibodies with different isotope labels were combined to produce the barcoded antibody mix.

### IMC protocol

FFPE sections were left at room temperature for 15 min after removal from storage. Deparaffinization was carried out using an AS-2 (Pathisto). The slides were placed into HIER buffer and heated at 95 °C for 30 min in a decloaking chamber (BioCare Medical) for epitope retrieval. Slides were cooled at room temperature for 20 min, washed in PBS for 15 min, and regions of interest were outlined with a hydrophobic pen (Vector Laboratories). Samples were then incubated with blocking buffer for 1 h at room temperature in a humidified chamber. Isotope-labeled antibody diluent was prepared in blocking buffer and incubated for overnight at 4 °C. Details of antibodies and their isotope labels are in Supplementary Table 1. Samples were washed in TBS for 15 min and incubated with 1:1000 dilution of 500 μM MaxPar Intercalator-Ir (Fluidigm) in PBS for 5-10 min, followed by a 15-min wash in TBS at room temperature. Slides were then dipped into deionized water for a few seconds, dried immediately using pressured air flow, and stored at room temperature until measurements. IMC images were acquired using a Hyperion Imaging System (Fluidigm). Laser ablation frequency was at 200 Hz, and pre-processing of the raw data into mcd files was completed using commercially available acquisition software (Fluidigm). Automated tuning of the argon flow and helium flow was performed on daily basis using a tuning slide coated with isotope-containing polymer (Fludigm).

### Data processing of IMC images into single-cell expression data

Steinbock v0.16.0 (https://github.com/BodenmillerGroup/steinbock) was used to convert the ion count raw data obtained from IMC software into single-cell expression data^16^. Briefly, DeepCell was used to segment the IMC images into single-cell masks. Aggregated images of cytoplasmic markers were used as cytoplasmic input, and iridium images were used as nuclei input. For CISI-IMC images, aggregated images of composite channels with cytoplasmic makers were used as cytoplasmic input. Single-cell expression data were obtained by calculating mean signal intensity for each channel in each cellular region defined by the cell mask. For the evaluation dataset, non-specific bright spots were observed in CD3 ground truth staining and to a lesser extend in other channels. Manual thresholding was applied to remove the cells with non-specific signal.

### SMAF algorithm

The dictionaries of protein expression modules were constructed from the training dataset using the SMAF algorithm. SMAF creates the dictionary by decomposing the training single-cell protein expression data (*X* ∈ ℝ ^*p×n*^) into a dictionary of protein expression modules (*U* ∈ ℝ^*p×d*^) and single-cell module activity (*W* ∈ ℝ^*d×n*^), where *p* is the number of proteins, *n* is the number of cells, and *d* is the number of modules in the dictionary. The steps of SMAF are as follows: (1) Initialize *U* and *W* by non-negative matrix factorization. The initial number of modules *d*_0_ = 80 was used for this work. (2) Fix *U* and calculate a sparse solution for *W* using Lasso or OMP. When using Lasso, the error tolerance coefficient *ldaW* is provided to calculate the sparsest *W* within the error tolerance defined by *ldaW*(= *λ*_*W*_) where 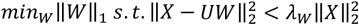. When using OMP, the sparsity *k* is provided to calculate the best fit solution, and up to *k* modules can be active (non-zero) for each cell *u*_*n*_ in *U* where 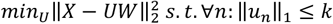.(2) Fix *W* and calculate the sparse solution for *U* using Lasso with the error tolerance coefficient of *ldaU*(= *λ*_*U*_) where 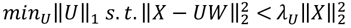. (3) Module-wise L2-normalization of *U*, that is, each module in *U* was scaled to the same vector size of 1. (4) Repeat steps (1), (2), and (3) for defined times. In this work, 100 iterations were used.

### CISI-IMC simulation protocol

We tested different dictionaries (*U* ∈ ℝ ^*p×d*^) and barcoding matrices (*A* ∈ ℝ ^*m×p*^) by simulating the CISI-IMC workflow using single-cell protein expression data (*X* ∈ ℝ ^*p×n*^) and evaluating the decompression performance, where *p* is the number of proteins, *d* is the number of modules in the dictionary, *m* is the number of composite channels, and *n* is the number of cells. CISI-IMC was simulated using the following steps: (1) Simulate single-cell composite data (*Y* ∈ ℝ ^*m×n*^) by compressing *X* using the barcoding matrix *A* by simple multiplication of *Y* = *AX*. (2) Calculate a sparse solution for *W* using Lasso. Note that the simulated composite data *Y* can be decomposed into *Y* = *AUW*, since *Y* = *AX* and *X* = *UW* according to the equations from compression and SMAF. As *A* and *U* are known, Lasso calculates the sparsest *W* within the error tolerance defined by *lda* (= *λ*) where 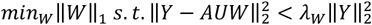. (3) Reconstruct decompressed single-cell protein expression data 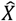 by calculation of 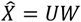. (4) Evaluate the decompression accuracy by calculating the correlation between the original *X* and the decompressed 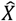. Pearson’s correlation coefficients were calculated separately for each protein and the mean and minimum of the correlations were typically used for evaluation, denoted as mean protein correlations and minimum protein correlations, respectively. The mean of correlations calculated separately for each cell was also used and are referred to as mean cell correlations.

### Normalizing single-cell protein expression data for simulating antibody titration

Only when simulating the antibody titration, we normalized *X* with weight *Wt*, denoted as 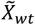, before simulating composite data. When weight is 1, protein-wise L2-normalization is performed, that is, each protein *x*_*p*_ in 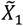 has the same vector size. When 0 < *Wt* < 1, the weighted mean of *X* and 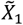 was calculated using the following equation: 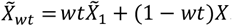.

### Finalizing the dictionary

Twelve SMAF parameter conditions were tested on the training dataset according to the CISI-IMC simulation protocol. For each condition, 200 randomly generated barcoding matrices were used. For general assessment of correlations between the original *X* and the decompressed 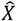, 25% of the training dataset was used for simulating CISI-IMC and the rest of the training dataset was used for SMAF dictionary calculation. The same strategy was used for assessing the performance on original *X* with weighted normalization. For testing the performance stability in different tissues in the training dataset, the CISI-IMC simulation was separately performed on data from each tissue, and the dataset excluding the tissue was used for SMAF dictionary calculation. Correlations between the original *X* and the decompressed 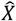 were calculated for each tissue and were averaged across tissues. Final SMAF parameters were selected based on the overall performance of general assessment, stability across tissues, and stability across different normalization weight for *X*. The finalized dictionary was obtained with the selected SMAF parameters (*algorithm*_*for*_*W*: *Lasso, ldaU*: 0.02, *ldaW*: 0.02, *Num*_*blocks*_*W*: 1) calculated on the entire training dataset.

### Finalizing the barcoding matrix

To finalize the barcoding matrix *A*, random *A*s barcoding 16 proteins in 8 composite channels were generated with the restriction that each protein maximally used 2 composite channels. The selection process was separated into two rounds. In the first round, decompression performance of 2000 randomly generated *A*s was evaluated on 25% of the training data according to the CISI-IMC simulation protocol. *U* was calculated using the finalized SMAF parameters on the rest of the training data. The simulation was performed four times, and the top 50 *A*s were selected from each simulation based on the minimum protein correlations, which resulted in 200 *A* candidates from 8000 *A*s tested. For the second round, another four simulations were performed, except that the 200 *A* candidates were fixed for each. Minimum protein correlations for each *A* were averaged across the simulations for the selection of the best *A*.

### CISI-IMC data acquisition and decompression using the evaluation dataset

The barcoded antibody mix for CISI-IMC was combined with the ground-truth antibody mix for the evaluation dataset. Co-staining of the ground-truth antibody mix provided the matched ground-truth *X* to compare against the decompressed *X*. Evaluation of actual CISI-IMC was performed as follows: (1) IMC data were obtained from tissues co-stained with the barcoded antibody mix and the ground-truth antibody mix. (2) IMC data analysis and single-cell segmentation yielded composite data *Y* and matched ground-truth *X*. (3) The sparse solution for *W* from *Y* was calculated using Lasso. Using known *A* and *U*, the sparsest *W* within the error tolerance defined by *lda* (= *λ*) where 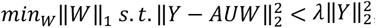, was determined using Lasso. (4) (skip to step 7 when not reweighting *A*) Reweight *A* with Lasso using known *W* and *U* . Lasso calculates the sparsest *Â* within the error tolerance defined by *ldaA*(= *λ*_*A*_) where 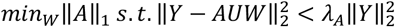. (5) Set all values to 0 in *Â* where original value of *A* was 0. (6) Repeat step 3 to 5, until the decomposition error (that is, 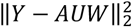) is minimized. In this work, we performed 6 iterations. Decompressed single-cell protein expression data 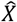 was reconstructed by 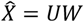. (7) Decompression accuracy was evaluated by comparing ground-truth *X* and decompressed 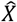.

## Data availability

Tiff files for IMC data and single cell data used in this manuscript will be made available on Zenodo upon acceptance.

## Code availability

All code used for preprocessing IMC data, SMAF dictionary calculation, and decompressing composite data are available at https://github.com/BodenmillerGroup/CISI-IMC_publication

## References

1. Schubert, W. et al. Analyzing proteome topology and function by automated multidimensional fluorescence microscopy. Nat. Biotechnol. 24, 1270–1278x (2006).

2. Gut, G., Herrmann, M. D. & Pelkmans, L. Multiplexed protein maps link subcellular organization to cellular states. Science (80-.). 361, (2018).

3. Lin, J. R., Fallahi-Sichani, M. & Sorger, P. K. Highly multiplexed imaging of single cells using a high-throughput cyclic immunofluorescence method. Nat. Commun. 6, 1–7 (2015).

4. Lin, J. R. et al. Highly multiplexed immunofluorescence imaging of human tissues and tumors using t-CyCIF and conventional optical microscopes. Elife 7, 1–46 (2018).

5. Gerdes, M. J. et al. Highly multiplexed single-cell analysis of formalinfixed, paraffin-embedded cancer tissue. Proc. Natl. Acad. Sci. U. S. A. 110, 11982–11987 (2013).

6. Goltsev, Y. et al. Deep Profiling of Mouse Splenic Architecture with CODEX Multiplexed Imaging. Cell 174, 968-981.e15 (2018).

7. Schürch, C. M. et al. Coordinated Cellular Neighborhoods Orchestrate Antitumoral Immunity at the Colorectal Cancer Invasive Front. Cell 182, 1341-1359.e19 (2020).

8. He, S. et al. High-plex imaging of RNA and proteins at subcellular resolution in fixed tissue by spatial molecular imaging. Nat. Biotechnol. 40, 1794–1806 (2022).

9. Ben-Chetrit, N. et al. Integration of whole transcriptome spatial profiling with protein markers. Nat. Biotechnol. 41, 788–793 (2023).

10. Liu, Y. et al. High-plex protein and whole transcriptome co-mapping at cellular resolution with spatial CITE-seq. Nat. Biotechnol. (2023) doi:10.1038/s41587-023-01676-0.

11. Giesen, C. et al. Highly multiplexed imaging of tumor tissues with subcellular resolution by mass cytometry. Nat. Methods 11, 417–422 (2014).

12. Keren, L. et al. MIBI-TOF: A multiplexed imaging platform relates cellular phenotypes and tissue structure. Sci. Adv. 5, 1–17 (2019).

13. Cleary, B., Cong, L., Cheung, A., Lander, E. S. & Regev, A. Efficient Generation of Transcriptomic Profiles by Random Composite Measurements. Cell 171, 1424-1436.e18 (2017).

14. Cleary, B. et al. Compressed sensing for highly efficient imaging transcriptomics. Nat. Biotechnol. 39, 936–942 (2021).

15. Ben-uri, R. et al. Escalating High-dimensional Imaging using Combinatorial Channel Multiplexing and Deep Learning. bioRxiv (2023).

16. Windhager, J., Bodenmiller, B. & Eling, N. An end-to-end workflow for multiplexed image processing and analysis. bioRxiv 2021.11.12.468357 (2021).

