## Supplementary Figures for "Compressed sensing expands the multiplexity of imaging mass cytometry"

### Extended Data Fig. 1

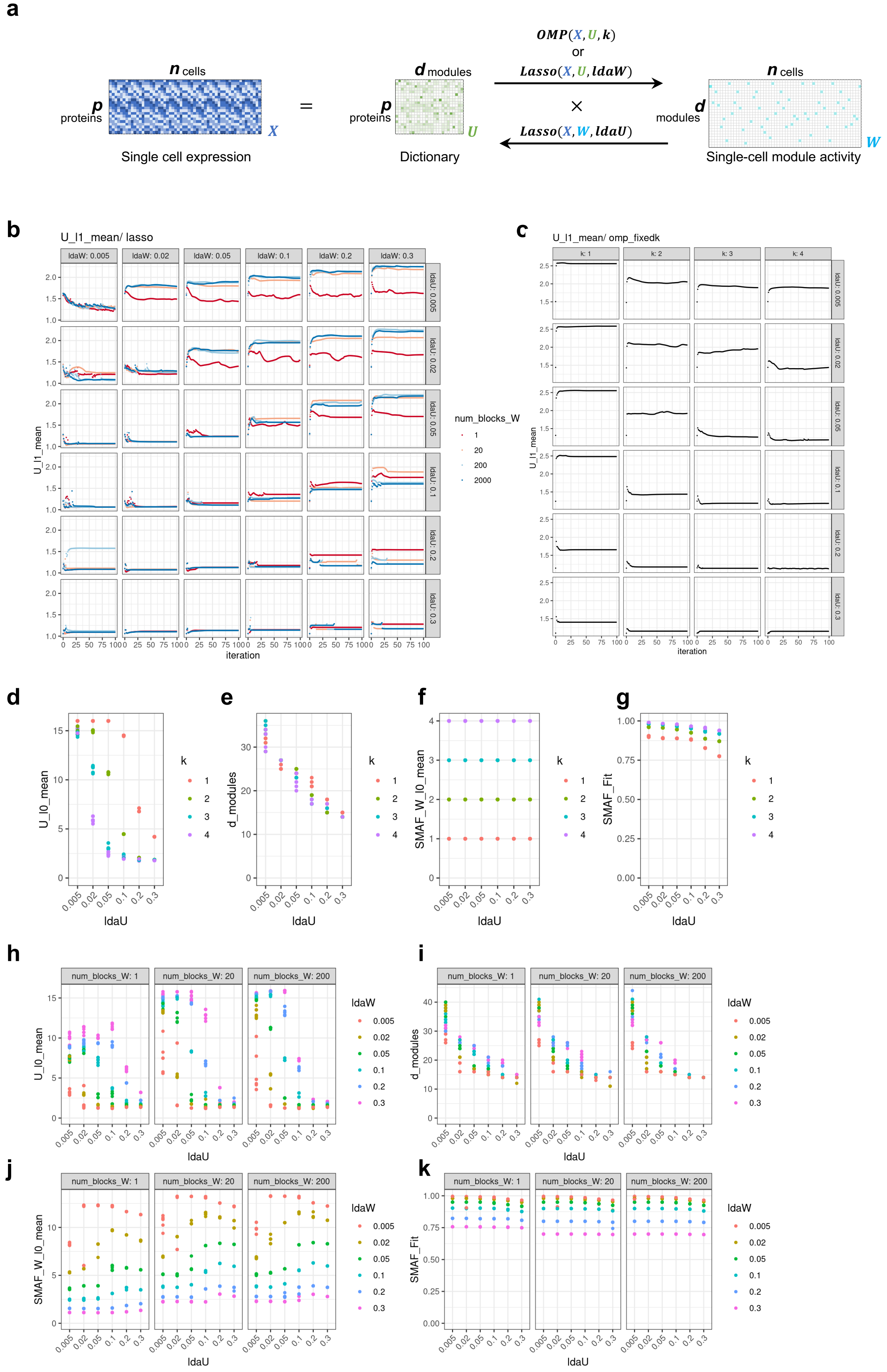

**Extended Data Fig. 1| SMAF algorithm optimization for CISI-IMC. a,** Schematic of SMAF algorithm workflow. SMAF decomposes single-cell protein expression data ($X$) into a dictionary of protein expression modules ($U$) and single-cell module activity ($W$) by iteratively finding the sparse solution for one while fixing the other. Lasso was used to calculate $U$ from a fixed $W$, and Lasso and OMP were used to calculate $W$ from a fixed $U$. $ldaU$ and $ldaW$ refer to the error tolerance coefficient used in Lasso when calculating $U$ and $W$, respectively. $k$ refers to the sparsity, the maximum number of modules each cell can express. **b-c,** Dictionary sparsity over 100 SMAF iterations calculated using **b**) Lasso or **c**) OMP. $U\_l1\_mean$: mean l1-norm (sum of absolute values in a module) across modules. Lasso enforces sparsity by minimizing the l1-norm. $ldaU$ and $ldaW$: error tolerance coefficients for Lasso. $Num\_blocks\_W$: Number of the groups of cells into which $W$ was separated when calculating $U$. The cells were grouped based on their vector size in $X$. $k$: Sparsity (number of non-zero input) per cell for OMP. **d-k,** Characteristics of $U$ and $W$ after 100 iterations of SMAF using **d-g**) OMP or **h-k**) Lasso to calculate $W$. Each parameter condition was tested 10 times. For panels d and h, sparsity of $U$ assessed by $U\_l0\_mean$, mean L0-norm (number of non-zero inputs) across modules. The lower L0 norm means the sparser $U$. For panels e and i, sparsity of $U$ assessed by the number of modules. For panels f and j, sparsity of $W$ assessed by $W\_l0\_mean$, mean l0-norm (number of non-zero inputs) across cells. The lower L0 norm means the sparser $W$. For panels g and k, accuracy of SMAF decomposition was assessed by the fit, subtracting the normalized distance between $X$ and $UW$ from 1; the closer the fit is to 1, the more accurate the decomposition was.

#### Extended Data Fig. 2

**a**

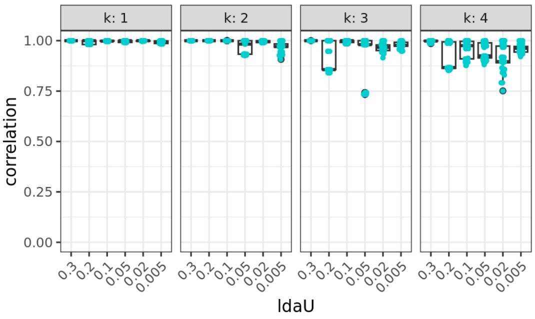


**b**

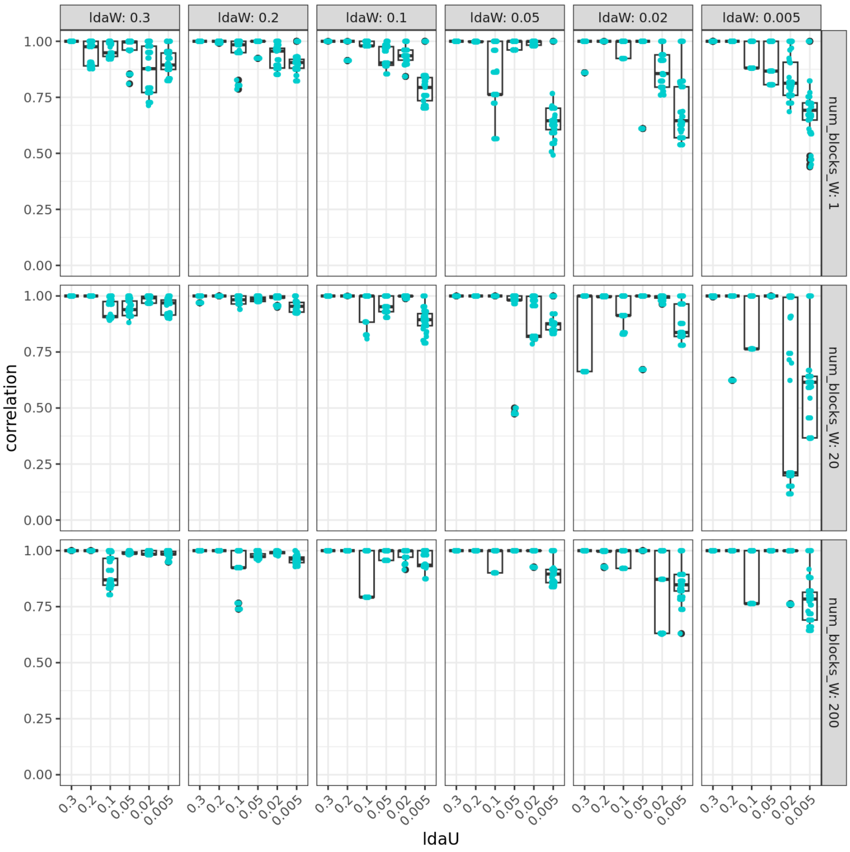


**Extended Data Fig. 2| SMAF stability over replicates. a-b,** Stability of SMAF algorithm assessed by the correlation of gene correlation matrix of $U$s between 10 replicates for each initial condition calculated using **a**) OMP or **b**) Lasso to calculate $W$. Correlations from the 45 possible comparisons within 10 replicates are plotted on top of the boxplot. The horizontal lines in the middles of boxes are the median values, and upper and lower boundaries of the boxes are 25th and 75th percentiles, respectively. Whiskers extend to maximum and minimum values within the 1.5 interquartile ranges.

#### Extended Data Fig. 3

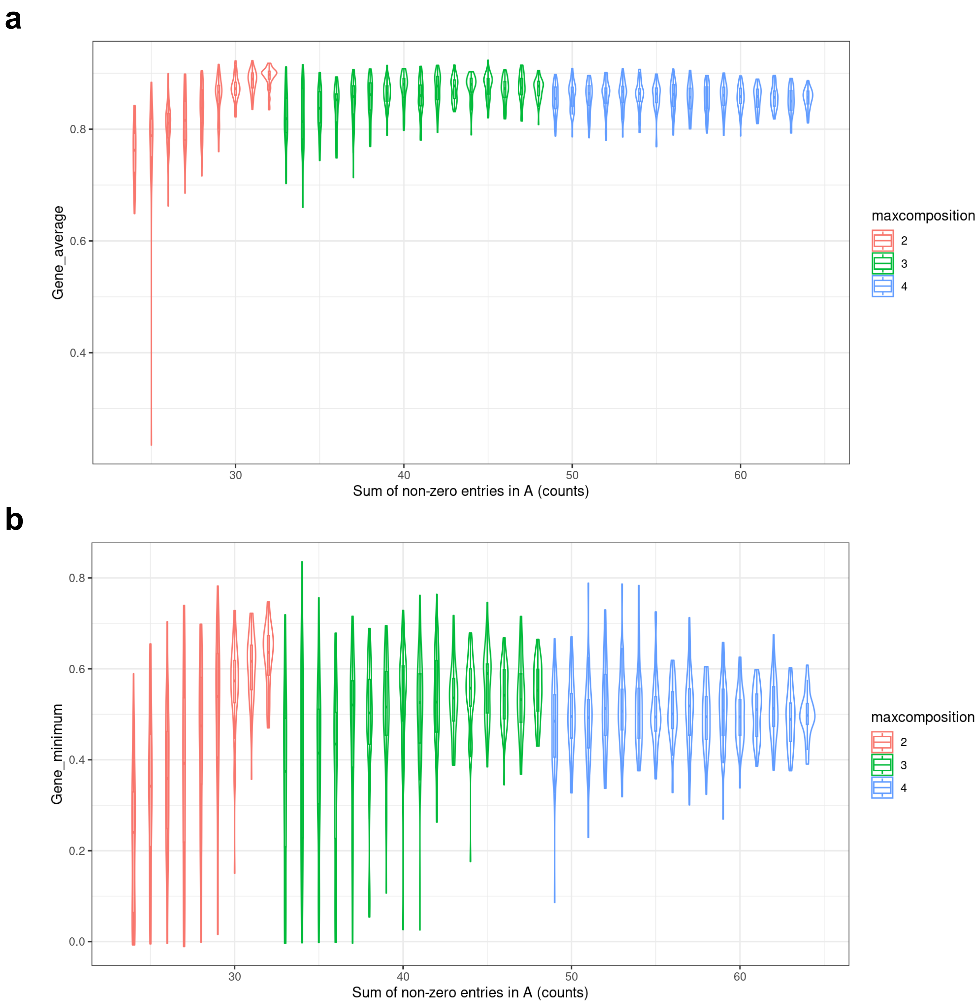


**Extended Data Fig. 3| Determination of maximum channels per protein in the barcoding matrix. a-b,** Violin plot overlayed by box plot for **a**) simulated protein-wise average correlation and **b**) simulated protein-wise minimum correlation between decompressed and original single-cell expression data determined using randomly generated barcoding matrices. Barcoding matrices were generated with the restriction of up to 2 (red), 3 (green), or 4 (blue) composite channels per protein, and 50 random barcoding matrices each for unique sum of non-zero entries were used for the simulation. The horizontal lines in the middles of boxes are the median values, and upper and lower boundaries of the boxes are 25th and 75th percentiles, respectively. Whiskers extend to maximum and minimum values within the 1.5 interquartile ranges.

### Extended Data Fig. 4

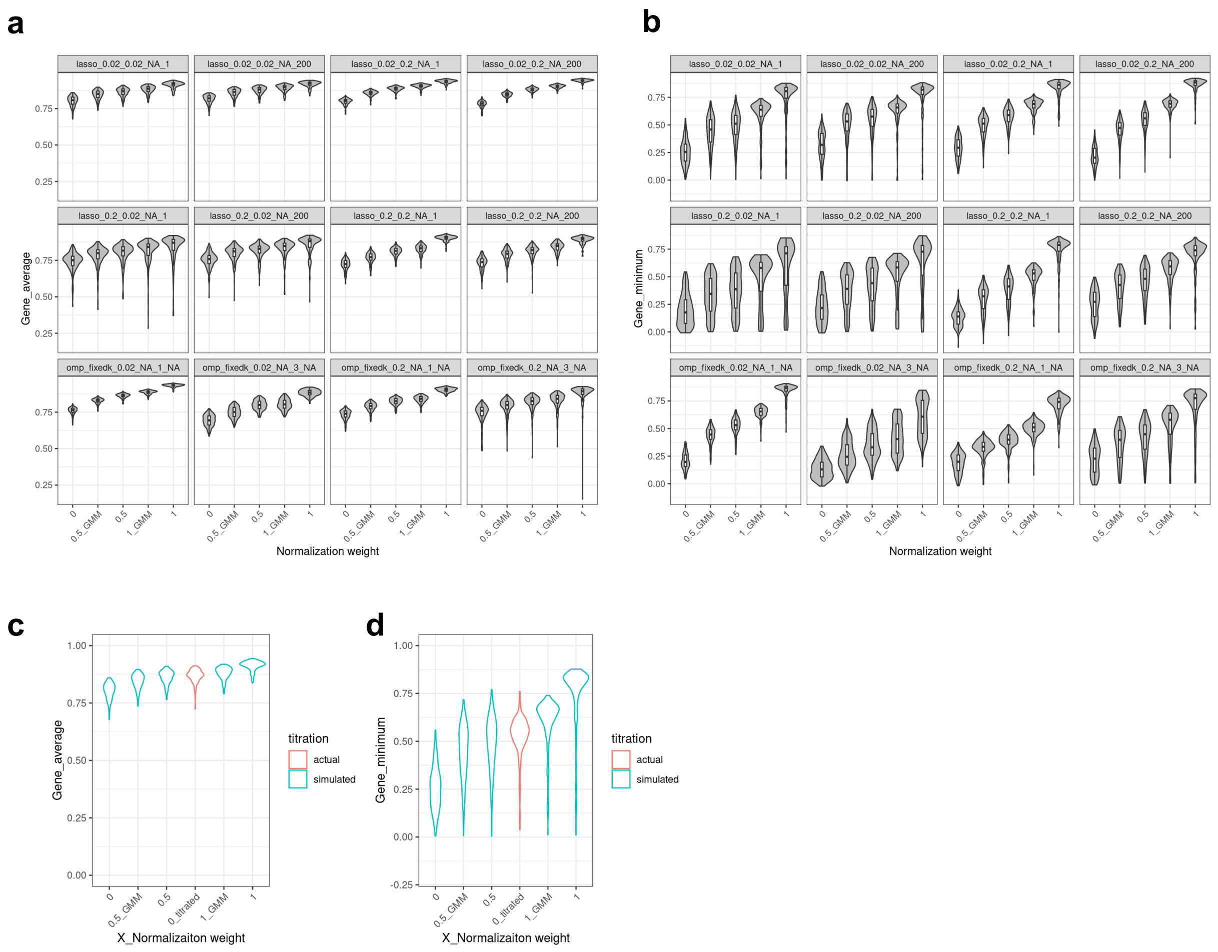


**Extended Data Fig. 4| Normalization of proteins from single-cell expression data. a-b,** Simulated **a**) protein-wise average correlation and **b**) protein-wise minimum correlation between decompressed and original single-cell expression data determined using 200 randomly generated barcoding matrices for each condition. The protein normalization strategies were the following: 0: no normalization; 0.5_GMM: mean of 1_GMM data and the data without normalization; 0.5: mean of l2-normalized data and the data without normalization; 1_GMM: normalization based on mean intensities of positive cell population defined by Gaussian mixture model; 1: l2-normalization (that is, vector size for each protein was the same after normalization). Twelve different SMAF parameter conditions were tested, labeled in the format of $(algorithm\_for\_W)\_(ldaU)\_(ldaW)\_(k)\_(Num\_blocks\_W)$. **c-d,** Simulated **c**) protein-wise average correlation and **d**) protein-wise minimum correlation between decompressed and original single-cell expression data determined using 200 randomly generated barcoding matrices for each condition. In addition to the normalization conditions used in panels a and b, single-cell expression data with manually titrated antibodies were used without normalization labeled as “0_titrated”.

### Extended Data Fig. 5

**a**

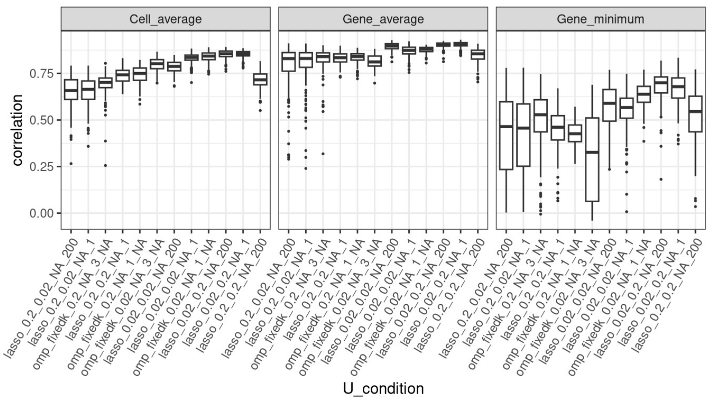


**b**

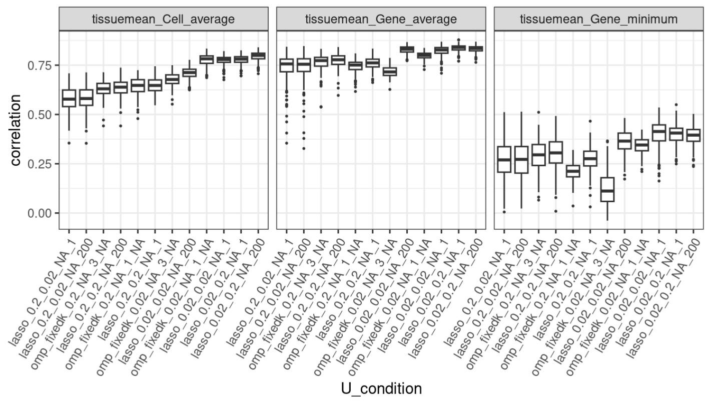


**c**

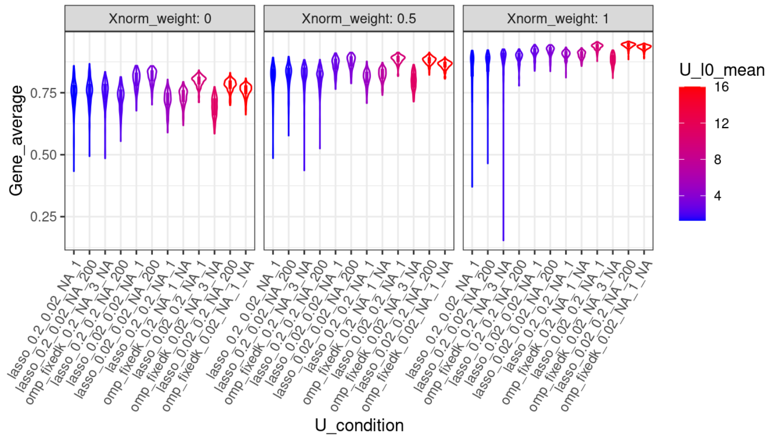


**d**

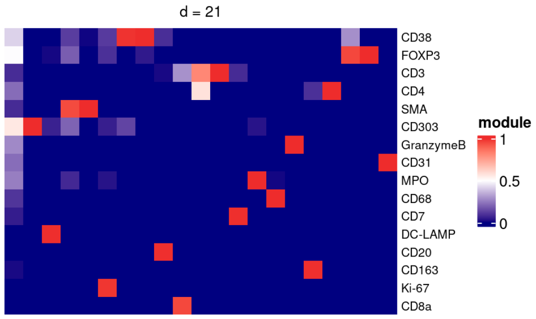

**Extended Data Fig. 5| Finalization of SMAF parameters. a,** Boxplot of simulated protein-wise average correlations (left), protein-wise minimum correlations (middle), and cell-wise average correlations (right) between decompressed and original single-cell expression data determined using randomly generated barcoding matrices. For each SMAF parameter condition, the correlations were calculated for each tissue using $U$ calculated on the training data excluding the tissue of interest. Correlation values were averaged across tissue conditions and are plotted for 200 random barcoding matrices used for each condition. The horizontal lines in the middles of boxes are median values, and upper and lower boundaries of the boxes are 25th and 75th percentiles, respectively. Whiskers extend to maximum and minimum values within the 1.5 interquartile ranges. Data points outside the range of whiskers are individually plotted. **b,** Boxplots of simulated protein-wise average correlations (left), protein-wise minimum correlations (middle), and cell-wise average correlations (right) between decompressed and original single-cell expression data determined using 200 randomly generated barcoding matrices. For each SMAF parameter condition, 25% of training data was used for simulation of decompression and the rest was used to calculate $U$. The horizontal lines in the middles of boxes are median values, and upper and lower boundaries of the boxes are 25th and 75th percentiles, respectively. Whiskers extend to maximum and minimum values within the 1.5 interquartile ranges. Data points outside the range of whiskers are individually plotted. **c,** Violin plots overlayed by boxplots for simulated protein-wise average correlations between decompressed and original single-cell expression data calculated using 200 randomly generated barcoding matrices with normalization of protein expression data. Normalization weights of 0 (left), 0.5 (middle), and 1 (right) were used. For each SMAF parameter condition, 25% of training data was used for simulation of decompression and the rest was used to calculate $U$. Violin plots and boxplots are colored by the sparsity of the dictionary $U$. The horizontal lines in the middles of boxes are the median values, and upper and lower boundaries of the boxes are 25th and 75th percentiles, respectively. Whiskers extend to maximum and minimum values within the 1.5 interquartile ranges. Data points outside the range of whiskers are individually plotted. **d,** Final dictionary $U$ obtained using all training single-cell protein expression data.

### Extended Data Fig. 6

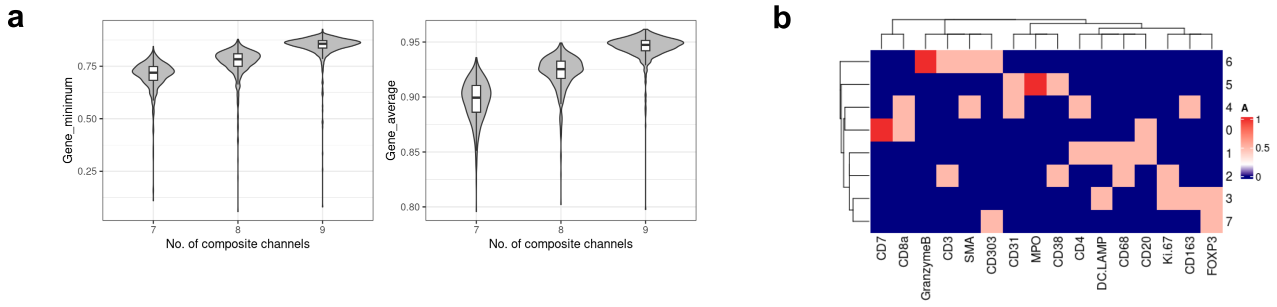


**Extended Data Fig. 6| Finalization of the barcoding matrix. a,** Violin plots overlayed by boxplots for simulated protein-wise minimum correlations (left) and protein-wise average correlations (right) between decompressed and original single-cell expression data calculated using top 200 barcoding matrices selected from 8,000 randomly generated barcoding matrices with 7, 8, or 9 composite channels for 16 proteins. The horizontal lines in middles of boxes are the median values, and upper and lower boundaries of the boxes are 25th and 75th percentiles, respectively. Whiskers extend to maximum and minimum values within the 1.5 interquartile ranges. **b,** Final barcoding matrix $A$ compressing 16 proteins into 8 composite channels.

**Extended Data Fig. 7**

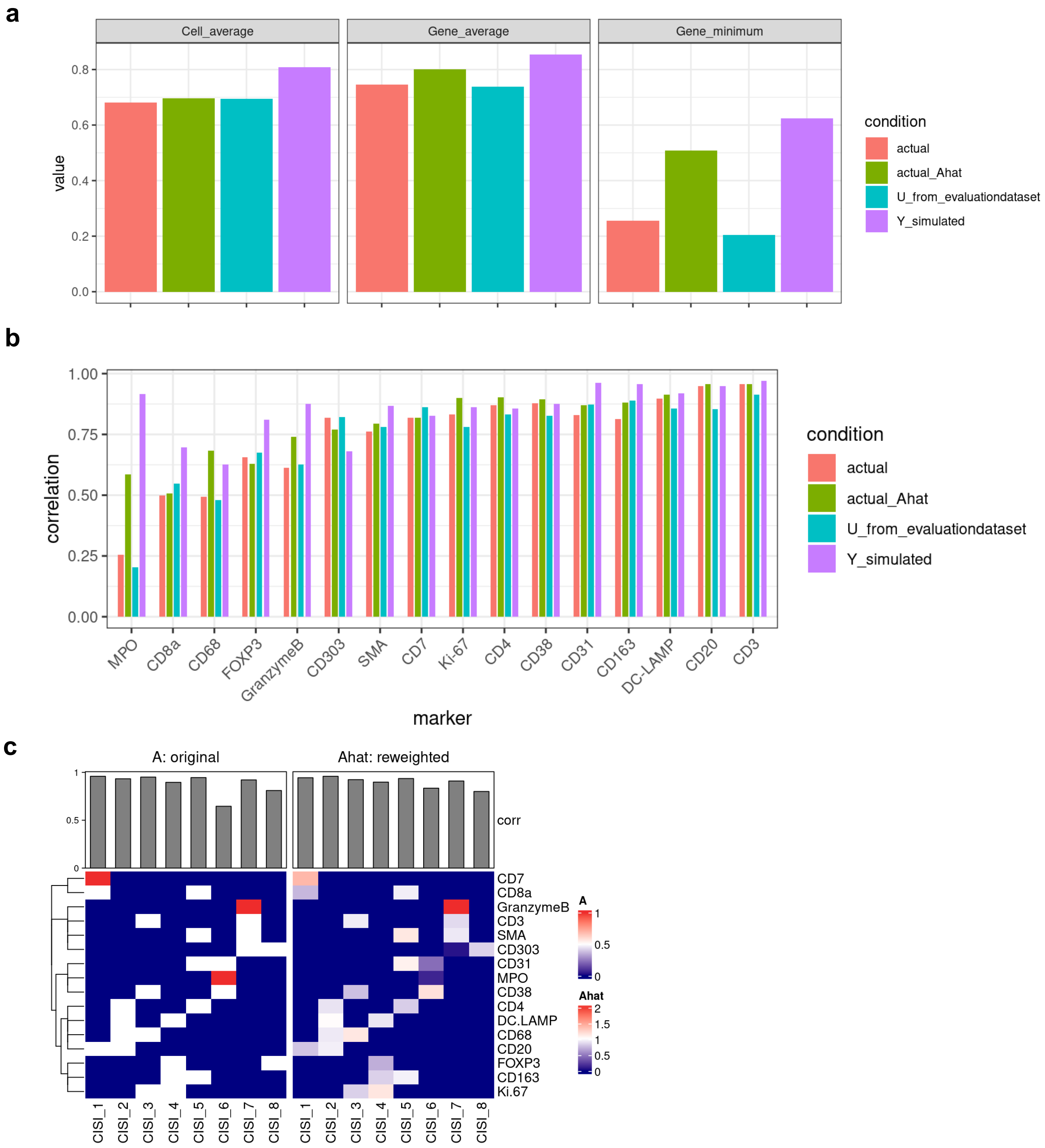


**Extended Data Fig. 7| Experimental analysis of CISI-IMC performance. a,** Cell-wise average correlations (left), protein-wise average correlations (middle), and protein-wise minimum correlations (right) between decompressed and original single-cell expression data. *Actual*: Decompression was performed using measured single-cell composite data and a dictionary calculated from single-cell protein expression data of the training dataset. *Actual_Ahat*: Decompression was performed using measured single-cell composite data and a dictionary calculated from single-cell protein expression data of the training dataset, while iteratively reweighting the barcoding matrix during the decompression. *U_from_evaluationdataset*: Decompression was performed using measured single-cell composite data and a dictionary calculated from ground-truth single-cell protein expression data. *Y_simulated*: Decompression was performed using simulated single-cell composite data using ground-truth single-cell protein expression data and a dictionary calculated from single-cell protein expression data from the training dataset. **b,** Correlations between decompressed and original single-cell expression data for each protein. **c,** Correlations between actual single-cell composite data and simulated composite data calculated using matching ground-truth single-cell protein expression data. Original barcoding matrix and reweighted barcoding matrix was used to simulate composite data. Correlations were calculated for each composite channel and are plotted with the barcoding matrix to show the proteins included in each composite channel.

#### Extended Data Fig. 8

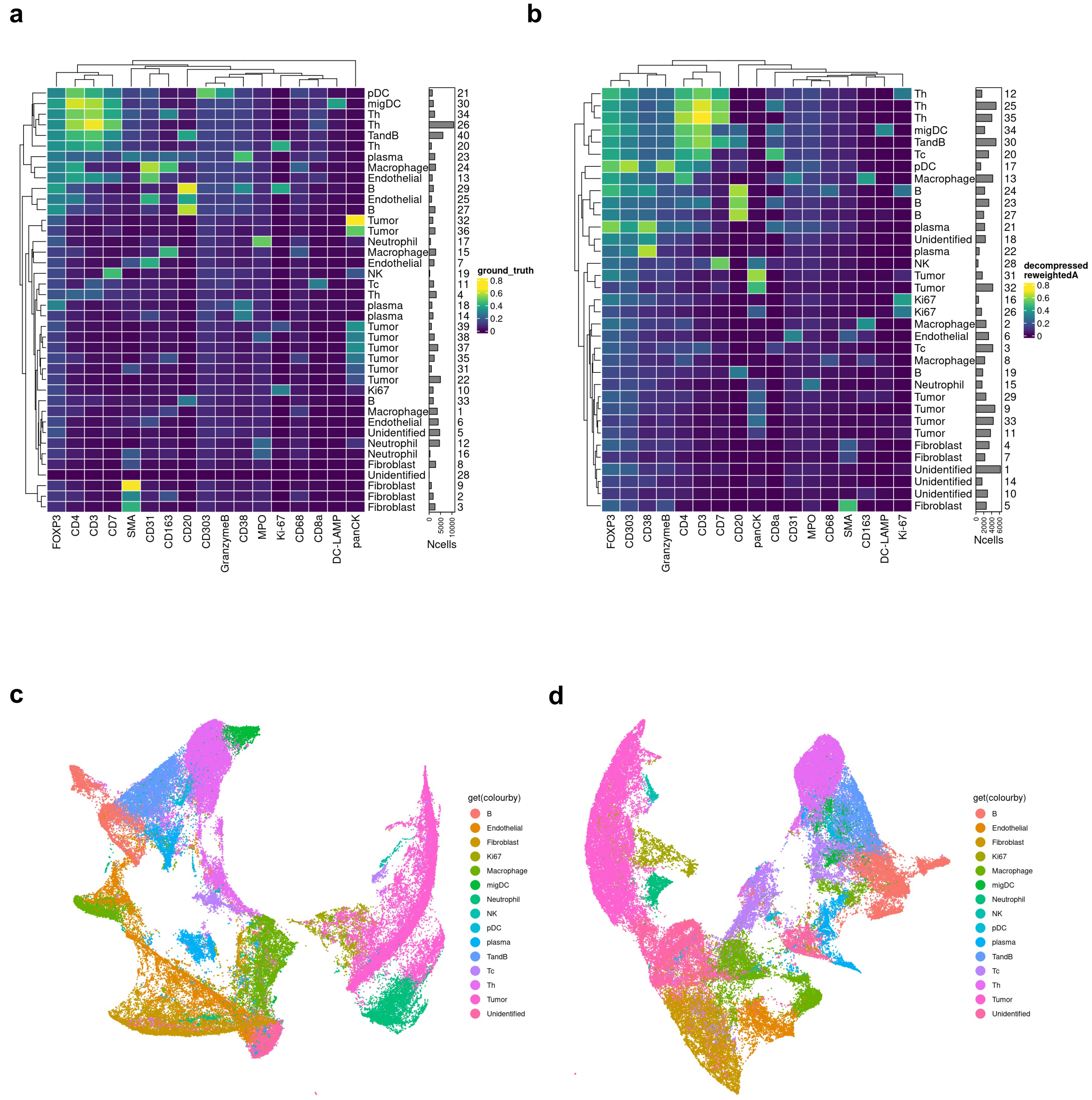


**Extended Data Fig. 8| Single-cell analysis of CISI-IMC decompressed data. a-b,** Phenograph clustering of **a**) ground truth and **b**) decompressed single-cell expression data labeled with annotated cell type and bar plots indicating the number of cells for each cluster. **c-d,** UMAP of **c**) ground truth and **d**) decompressed single-cell expression data, colored by the annotated cell type.
